# TARRAGON: Therapeutic Target Applicability Ranking and Retrieval-Augmented Generation Over Networks

**DOI:** 10.1101/2025.04.19.649662

**Authors:** Jon-Michael T. Beasley, Kara Schatz, Elvin Ding, Marcello DeLuca, Nahed Abu Zaid, Nyssa N. Tucker, Rada Y. Chirkova, Daniel J. Crona, Alexander Tropsha, Eugene N. Muratov

## Abstract

The identification of therapeutic protein targets is fundamental to the success of drug development and repurposing. Traditional approaches for target selection require extensive preclinical evaluation for toxicity and efficacy, making the process time-intensive and resource-heavy. Computational tools that efficiently prioritize and validate novel targets are needed to streamline drug discovery workflows. To address this gap, we developed TARRAGON: <u>T</u>herapeutic <u>T</u>arget <u>A</u>pplicability <u>R</u>anking and <u>R</u>etrieval-<u>A</u>ugmented <u>G</u>eneration <u>O</u>ver <u>N</u>etworks, a computational framework that integrates data mining and machine learning to identify, rank, and assess target-disease relationships to nominate new therapeutic targets. TARRAGON mines knowledge graphs to uncover meta-paths, or rules of graph traversal, linking potential therapeutic targets to diseases. It employs a classification model to rank target-disease hypotheses based on evidence patterns and utilizes a retrieval-augmented generation workflow to prompt a large language model for generating feasibility reports on prioritized targets. Using TARRAGON, we prioritized potential drug targets for non-muscle invasive urinary bladder cancer. Top-ranked candidates were validated using CRISPR gene effect and expression data from the Broad Institute DepMap portal. We further proposed chemical modulators for these targets to inform combination drug screening alongside approved bladder cancer therapeutics. TARRAGON introduces a novel, interpretable computational pipeline for therapeutic target discovery and pharmaceutical candidate nomination, offering the potential to accelerate drug development across diverse disease areas.

## Introduction

Of the approximately 30,000 human genes, only around 10% encode proteins considered “druggable” by small molecules or similar agents.^1^ Estimates suggest that 3,000^2^ to 10,000^3^ human genes are disease-related. The overlap between druggable protein-coding and disease-related genes is therefore estimated to comprise 600-1,500 human genes.^1–3^ Currently, over 850 human proteins are targeted by drugs with U.S. Food and Drug Administration (FDA) approval^4^, which has led to concerns that opportunities for novel target identification may be diminishing. However, scientific advancements over the past decade, including machine learning, artificial intelligence and advanced data analytics, have provided opportunities the scientific community to learn more about previously understudied proteins within known “druggable” protein families, allowing for the identification of novel targets. For instance, many of the over 500 kinases that comprise the “dark kinome” remain poorly characterized, but drug development groups have begun to explore these targets;^5,6^ development of inhibitors of the over 230 chromatin regulatory proteins has continued to expand; and the over 700 members of the ubiquitin-proteasome system are being actively investigated as potential drug targets.^4^ In parallel, recent advances in medicinal chemistry have opened the door for small molecule, peptidomimetic, or biologic drugs that target protein-protein interactions (PPIs), which were previously considered “undruggable” targets in drug discovery. In 2020 alone, 31 clinical trials were initiated to evaluate candidates targeting 9 unique PPI target systems.^7^ These trends showcase the opportunities for clinical translation arising from novel target identification. It is thus imperative that target selection workflows incorporate the latest knowledge of protein functions, PPIs, and pathway analyses.

Classical approaches to target discovery often rely on genetic manipulations of candidate targets using molecular biology techniques such as gene knockouts, RNA interference, or DNA mutagenesis, as well as population-level genome-wide association studies (GWAS). In some cases, observed phenotypic effects of small molecules are followed by deconvolution of those molecules’ targets using biochemical tools.^8^

However, there is growing focus in both academia and the pharmaceutical industry on developing computational methods to identify therapeutic targets and drug repurposing opportunities. These methods leverage prior discoveries and integrate diverse biomedical data to drive inference. One such approach is the biomedical knowledge graph (KG), a collection of entities or concepts (represented as nodes) and relationships between those entities (represented as edges). Nodes and edges are structured as “semantic triples”, or directional subject-predicate-object descriptions of knowledge that can be both readily processed by a machine or expressed in human natural language. KGs integrate disparate biomedical databases into a single unified machine-readable data structure that can be used to infer new knowledge through reasoning algorithms. This nascent approach will drive new knowledge of protein functions, PPIs, and pathway analyses, and will ultimately aid in identification of novel therapeutic targets.

Our team has remained at the vanguard of therapeutic target identification by developing a KG-based question-answering system called the Reasoning Over Biomedical Objects linked in Knowledge-Oriented Pathways (ROBOKOP).^9,10^ We have demonstrated its ability to uncover novel drug-disease associations by elucidating Clinical Outcome Pathways (COPs), which are mechanistic networks of biological entities that link known drugs and diseases in previously unexplored ways.^11^ We have also applied a KG mining approach to elucidate Adverse Outcome Pathways (AOPs)^12^ which identified the mechanisms of metal toxicity observed in patients experiencing complications from implanted medical devices.^13^

Building on the COP and AOP approaches, this work presents the development of a novel KG mining technology to discover Target Engagement Pathways (TEPs), which we call Therapeutic <u>T</u>arget <u>A</u>pplicability <u>R</u>anking and <u>R</u>etrieval-<u>A</u>ugmented <u>G</u>eneration <u>O</u>ver <u>N</u>etworks (TARRAGON). TARRAGON aims to uncover functional and mechanistic biological networks linking therapeutic targets to disease phenotypes. The approach involves three key steps: first, applying a rule-mining algorithm to extract explanatory TEP rules that highlight potential target-disease relationships; second, enhancing existing pathway-based methodologies (e.g., degree-weighted path count (DWPC) features)^14^ to train a classification model for link prediction and prioritize strong target-disease hypotheses; and third, implementing a retrieval-augmented generation (RAG) workflow using a large language model (LLM) to provide context-driven summaries of target feasibility based on TEP metapaths.

In this work, we demonstrate the utility of TARRAGON by applying it to nomination of targets for the treatment of urothelial bladder cancer (UBC). UBC is a common and deadly disease globally and represents a substantial burden to patients, providers, and health-systems. It is the 9^th^ most commonly diagnosed cancer worldwide with over 600,000 new diagnoses in 2022 alone.^15^ In the United States, there were nearly 17,000 bladder cancer-related deaths in 2024.^16^ While patients with metastatic bladder cancer have relied on platinum-based chemotherapy for more than four decades^17,18^, immune checkpoint inhibitors (e.g., pembrolizumab)^19,20^, the first targeted therapy called erdafitinib^21^, and an antibody–drug conjugate called enfortumab vedotin^19^ have been FDA approved in the past decade for the treatment of metastatic bladder cancer. Despite these therapeutic advances, metastatic bladder cancer remains essentially incurable, with a 5-year survival rate of only 8%.^22^ Thus, there is an unmet public health need to develop novel therapeutics that prolong overall survival for patients with urothelial bladder cancer. We applied the TARRAGON method to identify promising targets and drug candidates which could be tested for synergistic effects in combination with existing therapeutics.

## Results

### Development of a platform to obtain rules for target nomination

To begin the process of platform development, we curated a target-disease pair dataset for rule-mining. We queried the ROBOKOP KG to recover a total of 5,445 unique Target-Disease pairs. Of these, 2,714 of these pairs come from DrugCentral^26,27^, 1,958 come from DrugMechDB^25^, and 773 pairs were described in both primary knowledge sources. These Target-Disease pairs served as a ground-truth set of known therapeutic relationships used for rule mining and machine learning model training and validation. We then applied the AMIE 3 rule mining algorithm [cite] to delineate all rules with a maximum of 3 hops (i.e., 3 edges connecting Target and Disease nodes) associated with the defined “target_for” relationships. This process identified 81,361 rules linked to the “target_for” node pairs at least once. After converting rules to Cypher queries, removing duplicates, and applying statistical filters (See Methods), we recovered a total of 496 rules to evaluate for target nomination.

### A Target-Disease Ensemble Model for Virtual Screening

Following rule identification, the 496 rules were formulated as Cypher queries to retrieve all rule-defined paths connecting any combinations of the 1,351 Target and 813 Disease nodes. This process returned approximately 1 million possible Target-Disease pairs. Of these, 5,123 pairs corresponded to the 5,454 known “target_for” relationships, which we designated as positive examples for the training and evaluation datasets. The remaining 994,151 pairs were labeled as negatives, representing the space of false or unknown hypotheses potentially recovered through the application of these rules. Partitioning of training and validation sets resulted in 4,426 positive and 425,805 negative examples for training, while 697 positive and 569,043 negative examples were set aside for external validation.

### Development of a Target-Disease Ensemble Model for Virtual Screening

We developed an ensemble of 10 random forest models to classify Target-Disease pairs. Each model was trained with the full set of n positive examples from the training dataset, combined with a random sample of n×10 negative examples to maintain a 1:10 positive-to-negative ratio. The final prediction for each Target-Disease pair was obtained by averaging the prediction outputs across all 10 models, resulting in a score ranging from 0.0 to 1.0. To evaluate model performance, we applied the ensemble of models to make predictions for external validation set Target-Disease pairs. **Figure 3A** shows the model performance metrics for all predictions made on the 697 positive and 569,043 negative external data points. Importantly, the Target-Disease classifier model ensemble correctly predicted 86.5% of external positives and achieved a positive predictive value (PPV) of 0.584.

**Figure 1:**
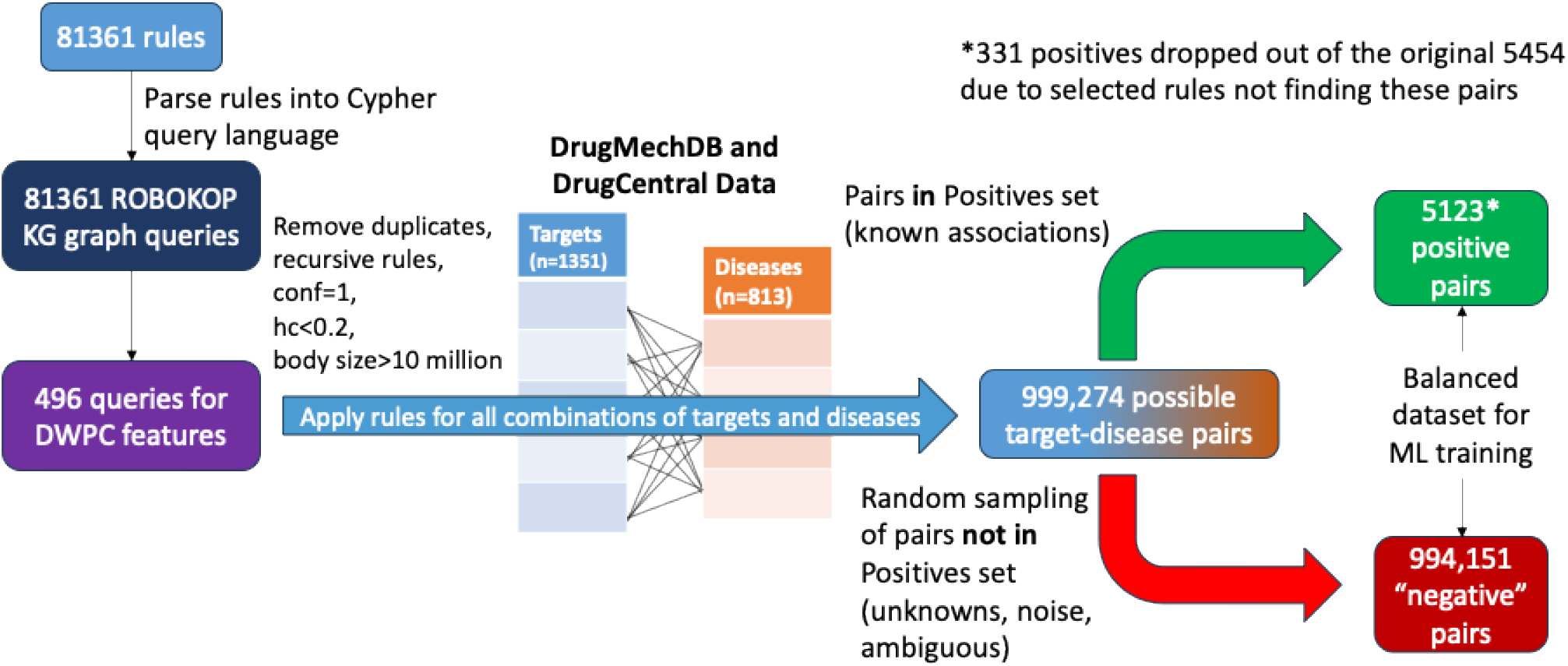
Rules discovered by AMIE 3 were converted to Neo4j Cypher queries for KG application. Rules were filtered to remove rules with any of the following features: duplicate, recursive (circular patterns), confidence=1, head coverage < 0.2, and body size > 10 million. 496 rule-based metapath queries were then applied to the KG to recover all possible paths between pairs of targets and diseases in the positive “target for” edge dataset. Positive and generated negative pairs were then used for model training and evaluation.

**Figure 2:**
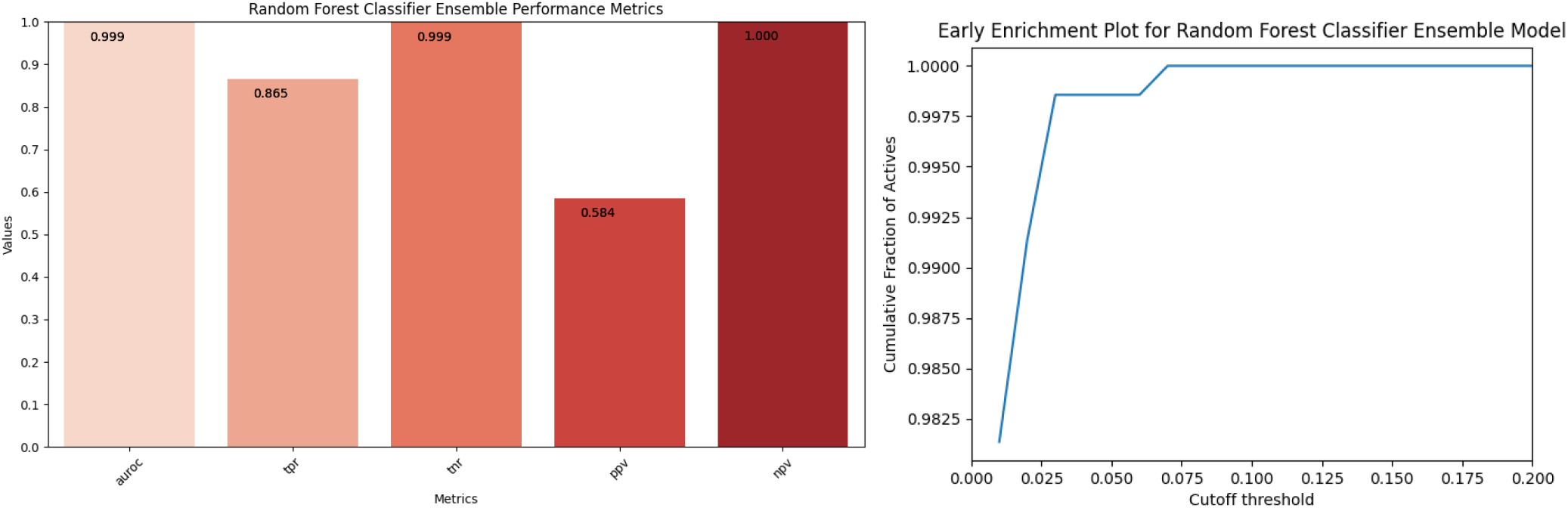
Random Forest ensemble performance on external set data. **A:** Area under ROC (AUROC), true positive rate (TPR), true negative rate (TNR), positive predictive value (PPV), and negative predictive value (NPV) for classification performance on the external dataset. **B:** Early enrichment curve shows that rank-ordered predictions on external data recover all positives within the top 7% of ranked hypotheses.

**Figure 3.**
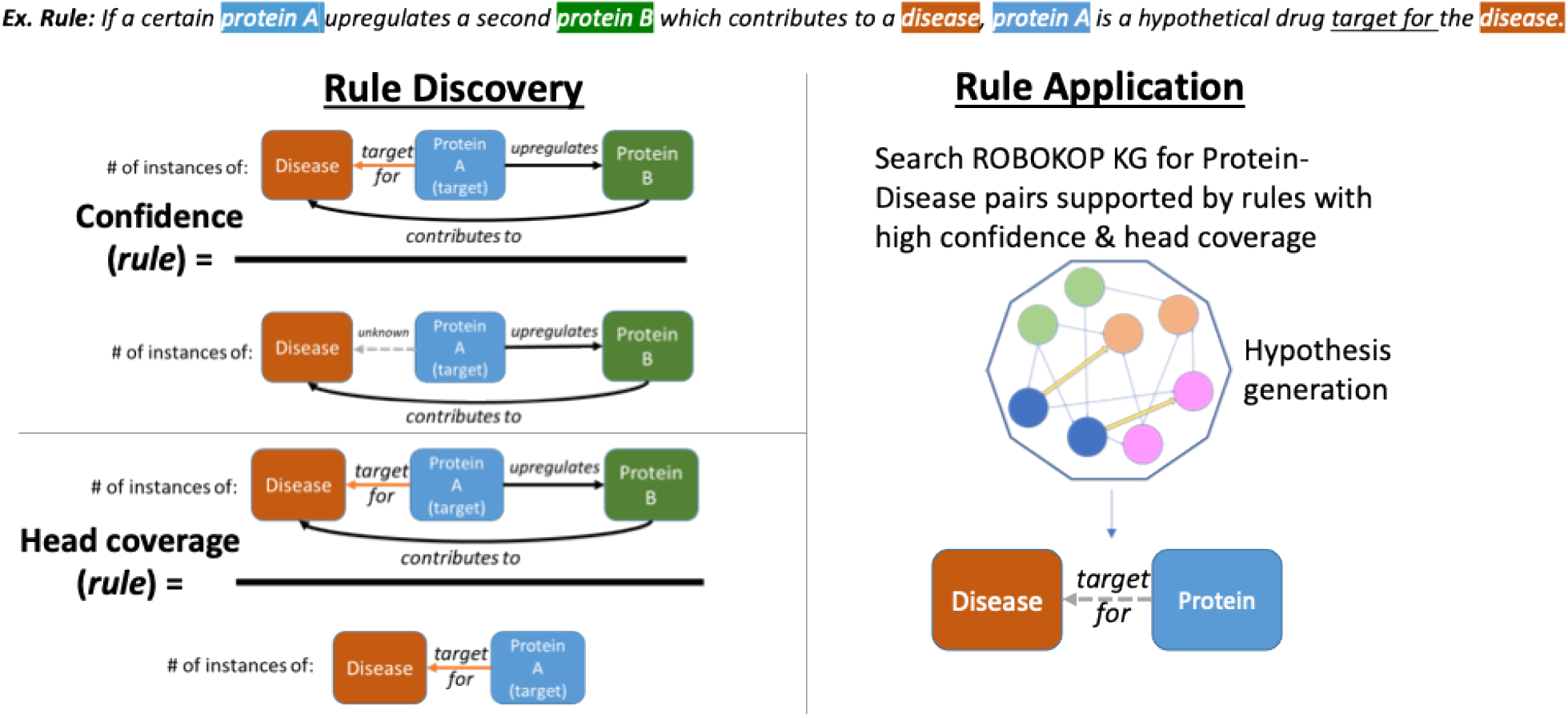
A description of rule mining with AMIE 3 and rule application on a KG. During the rule discovery stage, confidence, head coverage, and body size (denominator of confidence formula) for all patterns associated with a particular edge type (e.g., target for) are calculated. Rules are then converted to KG query format and applied to the graph to generate link prediction hypotheses.

To further assess the utility of the Target-Disease classifier output for hypothesis ranking, we calculated the enrichment factor of true positives found in the top *i-*th percentage of results when rank-ordering Target-Disease hypotheses by ensemble prediction output. All true positives fell within the first 7% of ranked hypotheses (**Figure 3B)**, suggesting the Target-Disease model has learned to generalize to external, unseen data, and correctly positions valid hypotheses within the top ranked answers.

### Virtual Screening for Bladder Cancer Targets

We next applied the Target-Disease ensemble model to a DWPC (W=0.2) dataset created for all gene or gene product nodes connected by any of the 496 rules to urinary bladder cancer to search for potential therapeutic targets. This process yielded 39,705 unique genes or gene products nodes. From these, we selected the top 100 human genes based on their mean positive class probability assigned by the Target-Disease random forest classifiers for further investigation (**Table 1**). For each of these top-ranked gene candidates, we queried the ROBOKOP KG to identify drugs that directly physically interact with the predicted targets. Our analysis specifically focused on FDA approved drugs that are either indicated for the treatment of bladder cancer or drugs/candidates that have been evaluated in bladder cancer clinical trials.

**Table 1:** Top 100 drug target candidates for urinary bladder cancer as predicted by the TARRAGON model. Green cells indicate known targets of FDA-approved drugs indicated to treat bladder cancer. Yellow cells denote targets of drugs involved in clinical trials for bladder cancer. White cells denote novel targets. * *TOP2A* was the only known target explicitly listed in the “target for” positive training dataset.

| Target Candidate Rank |  |  |  |  |  |  |  |  |  |
| --- | --- | --- | --- | --- | --- | --- | --- | --- | --- |
| 1-10 | 11-20 | 21-30 | 31-40 | 41-50 | 51-60 | 61-70 | 71-80 | 81-90 | 91-100 |
| TOP2A* | TUBB | TUBB6 | RRM2 | SP1 | MT-CO2 | MAPK3 | HDAC1 | CD274 | TP73 |
| TOP1MT | POLA1 | BAK1 | TUBB2B | FOLR2 | EGFR | CASP8 | PTGS1 | NOX1 | ADRA1B |
| TOP2B | PARP2 | CMPK1 | PTGS2 | TNKS2 | MDM2 | FLT1 | BAX | PDCD1 | H2BFS |
| TOP1 | TUBB8 | TUBA1A | SYT6 | ATIC | DHFR | TNF | PARP10 | PARP12 | PSMB1 |
| RARA | ABCB1 | PRKCD | CDC6 | PARP3 | AR | PTGES3 | IL32 | LLPH | ANKRD1 |
| POLE | RRM1 | TUBE1 | CYP1A2 | TUBG1 | CASP3 | YWHAB | SLCO2B1 | FNTB | ACHE |
| TUBB4B | TUBB1 | TNKS | GSR | TUBD1 | GART | CYP2C9 | SLC46A1 | L1RE1 | GSK3A |
| PARP1 | HIF1A | HMGB1 | UMPS | RIOK1 | NR3C1 | GTPBP10 | ATM | SLC47A1 | FLT4 |
| POLD1 | TUBB2A | PRP4K | TUBB4A | FASLG | CYP3A4 | TP53 | NR1I2 | CASP7 | GSTP1 |
| ICOSLG | TUBB3 | TYMS | GSK3B | SLAMF7 | MAPK12 | TNFRSF8 | ROS1 | RELA | H2AC12 |

We next scanned Pharos^33^ and ChEMBL^34^ for drug candidates which act on the remaining predicted targets. To select candidates with maximum repurposing potential, we focused the search to include previously FDA approved therapeutics, when possible, but also included high-potency investigational compounds when no approved therapeutics existed. These compounds, listed in **Table 2**, represent promising candidates to experimentally screen using bladder cancer cell lines and patient-derived organoids.

**Table 2:** Approved or investigational drug repurposing candidates were selected for the predicted targets which were not involved with an indication or clinical trial for bladder cancer. We applied a preference for approved drugs and compounds with high potency at the target. Target Central Resource Database (TCRD) target designation^33^ of each gene is listed next to gene symbols in parentheses. FDA approval status or max phase of clinical trials is listed next to drug names in parentheses.

| Target Name | FDA-Approved or Investigational Drugs | Drug Activity at Target |
| --- | --- | --- |
| RARA (Tclin) | Tretinoin (approved) | Agonist, pEC50: 6.7 |
|  | Bexarotene (approved) | Agonist, pEC50: 8.22 |
|  | Alitretinoin (approved) | Agonist, pEC50: 6.52 |
|  | Adapalene (approved) | Antagonist, pKi: 5.96 |
|  | Tazarotene (approved) | Agonist, pEC50: 7.2 |
|  | Trifarotene (approved) | Agonist, pEC50: 6.3 |
|  | Acitretin (approved) | Agonist |
|  | Etretinate (approved) | Agonist |
| ICOSLG (Tbio) | n/a | n/a |
| BAK1 (Tbio) | n/a | n/a |
| SYT6 (Tbio) | n/a | n/a |
| CDC6 (Tbio) | n/a | n/a |
| GSR (Tclin) | Carmustine (approved) | Inhibitor, pIC50: 5.09 |
| SP1 (Tbio) | n/a | n/a |
| FASLG (Tbio) | n/a | n/a |
| SLAMF7 (Tclin) | Elotuzumab (approved) | Antibody binding, pKd: 8.05 |
| MDM2 (Tchem) | Apomorphine (approved) | pIC50: 6.71 |
|  | Thiram (approved) | pEC50: 6.75 |
|  | Cytarabine (approved) | pIC50: 7.7 |
|  | Navtemadlin (phase 2) | pKd: 10.42 |
|  | Idasanutlin (phase 3) | pKd: 9.82, pIC50: 8.22 |
| CASP3 (Tchem) | CHEMBL359598 (preclinical) | pIC50: 10.96 |
| GART (Tclin) | Pemetrexed (approved) | Inhibitor, pKi: 6.42 |
|  | Lometrexol (phase 2) | pKi: 7.22 |
|  | CHEMBL82261 (preclinical) | pKi: 8.70 |
| CASP8 (Tchem) | CHEMBL443565 (preclinical) | pIC50: 10.80 |
| PTGES3 (Tbio) | n/a | n/a |
| YWHAB (Tbio) | n/a | n/a |
| GTPBP10 (Tbio) | n/a | n/a |
| TP53 (Tchem) | CHEMBL4248974 (preclinical) | pEC50: 8.7 |
| TNFRSF8 (Tclin) | Brentuximab Vedotin (approved) | Binding agent |
| BAX (Tchem) | CHEMBL4579601 (preclinical) | pIC50: 7.37 |
|  | CHEMBL4794290 (preclinical) | pEC50: 6.41 |
| IL32 (Tbio) | n/a | n/a |
| ATM (Tchem) | Dactolisib (phase 3) | pIC50: 8.15 |
|  | AZD-1390 (phase 1) | pIC50: 10.05 |
| NOX1 (Tchem) | Phenothiazine (phase 2) | pIC50: 6.46 |
| LLPH (Tbio) | n/a | n/a |
| L1RE1 (Tbio) | n/a | n/a |
| CASP7 (Tchem) | CHEMBL178116 (preclinical) | pIC50: 11.00 |
| TP73 (Tbio) | n/a | n/a |
| H2BFS (Tbio) | n/a | n/a |
| HIST1H2AH (Tbio) | n/a | n/a |

### Target Prioritization

Next, we analyzed the 29 predicted targets lacking a bladder cancer indication or clinical data by using the Broad Institute’s DepMap Data Explorer.^35,36^ We profiled the mean mRNA transcript expression (log2(TPM + 1)) across data available from 33 bladder cancer cell lines, and the mean CRISPR knockout gene effect on cell population (Chronos score^37^) across 30 bladder cancer cell lines for the 29 genes to assess the expected dependency of bladder cancer cell lines on the predicted targets. To prioritize the most promising candidate genes, we applied thresholds of mean expression > 4 log2(TPM + 1) and a mean CRISPR gene effect < -0.4 Chronos score. After applying these criteria, four target candidates remained: *CDC6, MDM2, GART*, and *LLPH*. These results are visualized in **Figure 4**, highlighting the significant dependencies of bladder cancer cells on these selected targets.

**Figure 4.**
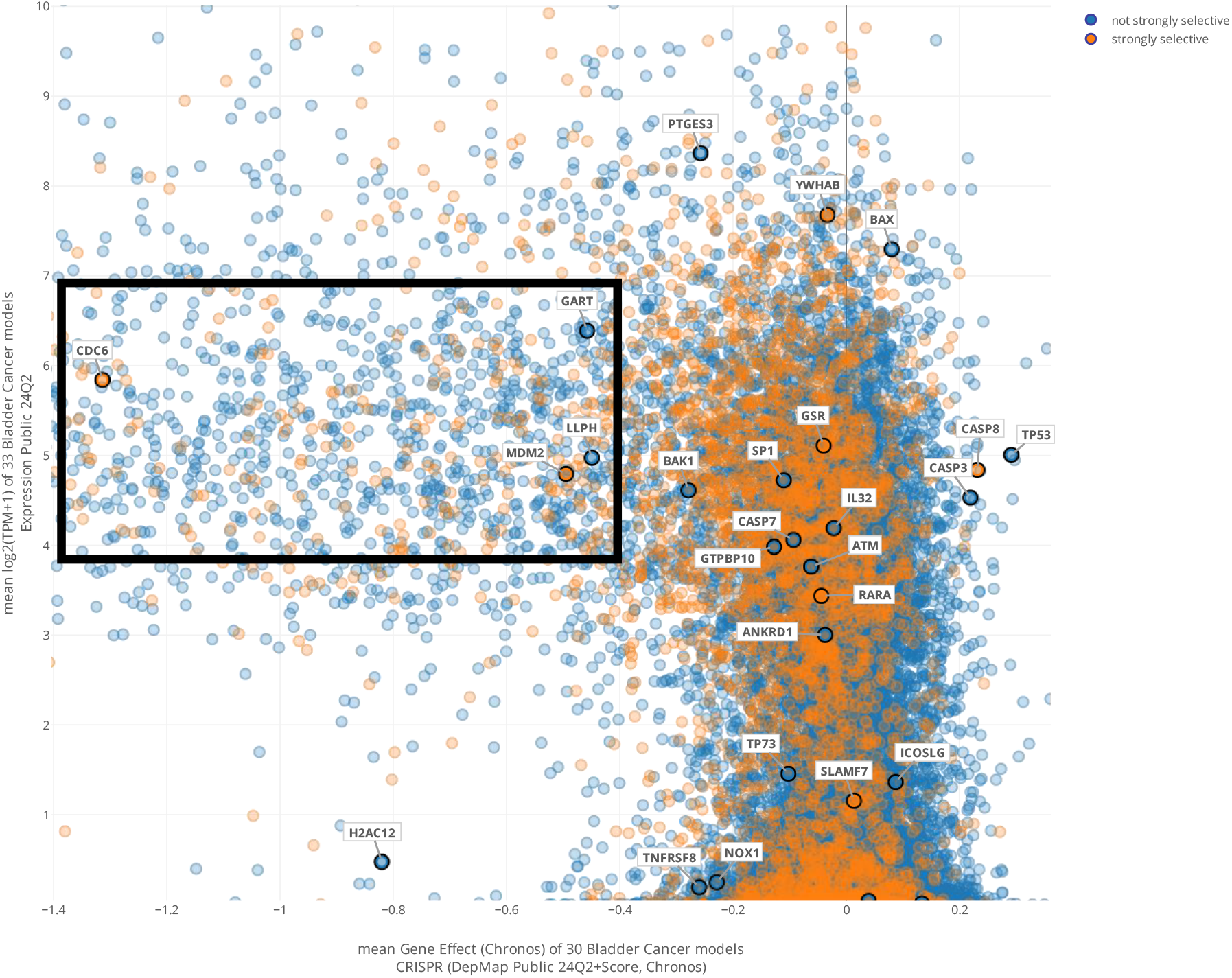
Mean dependence of various bladder cancer cell lines on individual candidate genes (CRISPR knockout Chronos score) was plotted against mean expression (log2(TPM) to highlight the candidates with relatively high expression that play an essential role in bladder cancer survival and proliferation. Of all novel target candidates, those shown in the black box (CDC6, MDM2, GART, LLPH) best fit the dependency and expression criteria for further consideration.

### Retrieval-Augmented Generation of Target Feasibility Reports

Because expert review of top Target-Disease hypothesis predictions is a critical part of experimental planning, we employed a RAG summarization workflow that supplements a general LLM prompt about the feasibility of predicted therapeutic target with background knowledge and context derived from KG paths extracted from TEP rule features used to make the predictions. We created LLM (GPT-4o) generated reports about the role of four putative targets of urinary bladder cancer (*CDC6, MDM2, GART, and LLPH*) to summarize the knowledge derived from the KG. Please note that the following texts, reported in **Supplemental Data**, are reported as-is from LLM-generation, and any inaccurate claims or citations are the product of LLM hallucination:^38^

## Discussion

This work provides a first-of-its-kind framework for KG-driven therapeutic target discovery using automated rule mining and metapath structural features to predict novel putative targets for diverse disease states. While previous works have elaborated on metapath-based link prediction and feature engineering for applications to gene-disease relationship prediction^14^ and prediction of novel Alzheimer’s disease-related genes^31^, these methods relied on the manual selection of explanatory metapaths, which is subject to human bias. Our method instead performs automated metapath discovery via graph rule mining^28^, which enables extensibility of our predictive modeling framework to future KG updates and expansions. Furthermore, statistical features of automatically mined rule allow data-driven selection and prioritization of rule-based metapaths used for model building.

Another recent work focused on target-disease prediction over KGs demonstrated the power of graph embeddings for predicting therapeutic targets.^39^ However, graph embedding-based models are considered “black box” models that provide little interpretability about how or why a predicted target was selected. While these embedding-based predictive models achieved high accuracy, little explanatory power can be derived from embedding features alone.

The TARRAGON model, on the other hand, enables prediction interpretability as each dimension of a target-disease pair representation vector corresponds to a TEP pattern used to retrieve specific answers. We take advantage of this feature by aggregating KG answer paths derived from high-DWPC metapaths and summarizing this knowledge in a concise, LLM-generated target feasibility report. Thus, the TARRAGON system stands apart from existing methods by providing easily digestible explanations of predictions derived from the same features used to make the predictions.

We evaluated performance of the TARRAGON prediction model on an external set specifically chosen so that no target or disease nodes overlapped with training data. This is an important criterion for evaluating performance on pairwise datasets such as the type considered in this work. Additionally, we included many external negatives (*n=*569,043) relative to external positives (*n=*697) to reflect the reality that applying 3-hop metapaths to our KG is expected to generate many, many more false hypotheses than true ones for any given disease. While the true positive rate (TPR) of 86.5% was quite promising, PPV was lower than ideal. Considering the extremely large class imbalance of the held-out data (1:816 positive:negative ratio), and the possibility that the synthetically generated negatives possibly contain some unknown, but viable, Target-Disease hypotheses or known facts missing from our “target for” dataset, this 0.584 PPV is an acceptable measurement.

Some limitations exist for TARRAGON and other metapath-based predictive models. First, the search space for automated rule mining can be exceedingly large considering that the number of *k*-hop linear paths for a graph with *R* relationship types is *Rk* and the AMIE 3 procedure must recover all unique instances of each rule. To overcome this challenge for mining *3*-hop rules, we removed the most abundant and least meaningful relationships from the rule mining graph and set the body size cutoff to 10 million unique instances of each rule. Similar considerations will need to be taken when applying the TARRAGON framework to other large biomedical KGs and may require some upfront intelligent selection of rule mining criteria. Second, application of rule-based metapaths to generate model training, test sets, and virtual screening sets for specific diseases follows an *O(m)* computational runtime that scales linearly with the number of metapaths *(m)* included in the feature space. The feature space can be reduced rationally by filtering rules by confidence, head coverage, and body size to achieve a reasonable run time. Additionally, datasets for virtual screening must be produced *de novo* to obtain target predictions for any specific disease or set of diseases, requiring a time-intensive dataset generation step prior to generating predictions. Compared to graph embeddings, which can be precomputed and rapidly accessed from a vector store, metapath-based approaches will always suffer from this limitation. Finally, as with any KG-based predictive model, the performance and applicability of the TARRAGON system is highly sensitive to structure and content of the KG being operated on. The DWPC featurization accounts for the inherent bias of metapaths to traverse high-degree nodes, but careful curation and data modeling standards for KG creation are essential to avoid introduction of other biases or erroneous statements. In this current work, we applied TARRAGON to an enhanced version of the ROBOKOP KG by supplementing the baseline graph with clinical trials (clinical-trials-kp)^40^, cell signaling events (SIGNOR 3.0)^41^, and kinase-substrate interactions (KinAce)^42^, while also removing any possibly erroneous relationships that resulted from text-mining. We envision that further enrichment and curation of a target-centric KG will continue to improve TARRAGON performance.

With a case study on urinary bladder cancer, we demonstrated the utility of TARRAGON as a framework for therapeutic target-disease link prediction and evidence summarization. Among the top 100 target candidates, 43 were targets of drugs indicated for bladder cancer, 28 were targets of drugs involved in clinical trials for the disease, and 29 represented novel therapeutic target candidates. Since only TOP2A was explicitly listed as a target for bladder cancer in the KG, the enrichment of so many other known and investigated bladder cancer targets in the top predictions showcases the power of TARRAGON for link prediction and KG completion under assumptions of incomplete evidence.

The novel targets predicted in this set motivate the design of biochemical and cell-based experiments involving known chemical inhibitors of these targets (**Table 2**) or arrayed CRISPR knockout/RNA silencing screens in specific bladder cancer cell lines to confirm the data from pooled CRISPR screening shown in **Figure 4**. Computational workflows for drug discovery should be paired with experimental validation to reflect their true utility, therefore we envision the next steps for TARRAGON development and evaluation to include collaboration with experimental scientists interested in applying our predictive system to real-world drug discovery problems.

## Methods

### Metapath-Optimized Knowledge Graph

The hypotheses generated by the TARRAGON system are derived from TEP metapaths within the extensive biomedical KG, ROBOKOP KG.^9^ To tailor it to our specific needs, we created a customized version of ROBOKOP KG using the ORION KG build pipeline.^23^ We used the ORION pipeline to generate the customized ROBOKOP KG as a Neo4j (v5.22.0) database, enabling us to process Cypher queries efficiently throughout the TARRAGON workflow. By carefully selecting high-quality data sources and refining the graph structure, we ensured that the customized ROBOKOP KG provided a robust and reliable foundation for extracting and processing Target-Disease hypotheses.

The data sources included in this customized KG were defined in the graph_spec.yml build specifications file and included: BINDING-DB, CHEBIProps, ClinicalTrialsKP, Comparative Toxicogenomics Database (CTD), DrugCentral, DrugMechDB, GtoPdb, GWASCatalog, Hetio, HGNC, HMDB, HumanGOA, IntAct, KinAce, MonarchKG, MONDOProps, Ontological Hierarchy, PANTHER, PHAROS, Reactome, SIGNOR, STRING-DB-Human, and UbergraphNonredundant.

To improve the accuracy and efficiency of the KG, we excluded certain sources and edge predicates that posed challenges. Specifically, we omitted the “textminingkp” knowledge source^24^, as it contains assertions automatically mined from publication texts, which are often prone to inaccuracies and mistakes resulting from the language-model which generates the assertions. Additionally, we removed the “biolink:related_to” and “biolink:subclass_of” predicates due to their ambiguity and excessive abundance, which could negatively impact rule mining, both in complexity/runtime and accuracy of rules extracted from this process.

### Selection of Target-Disease Pairs

We defined a “Target-Disease pair” as a protein target and its associated disease, where an approved drug is indicated for the disease and exerts its therapeutic action to treat that disease *via* modulation of the target. We extracted Target-Disease pairs from the DrugMechDB^25^ and DrugCentral^26,27^ using the ROBOKOP KG by constructing a Cypher query:

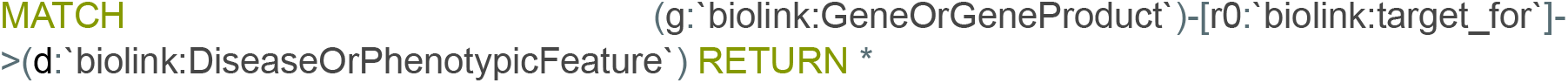

### Mining and Selection of Target-Disease Association Rules

We define KG rules as sequences of directed edge labels, capturing linear, *k*-hop evidence patterns between source and target nodes linked by the specified “target for” edge label. We employed the Association Rule Mining under Incomplete Evidence (AMIE 3) algorithm^28^ to efficiently mine TEP rules, or metapaths, of KG traversal statistically associated with known examples of “target for” semantic triples. AMIE 3 calculates key statistical metrics, including head coverage, confidence, and body size of each rule, allowing filtering of rules by statistical features. We converted each of these rules into Neo4j Cypher queries which could be applied to mine potential drug target hypotheses from the ROBOKOP KG. We further process these rules by removing duplicates (rules which recover the same answers) and recursive rules. We also remove rules with confidence = 1 to prevent data leakage when the presence of a certain feature only correlates with the positive class^29^, rules with head coverage < 0.2 to prioritize recall of positives, and rules with body size > 10 million to improve rule application speed and minimize extraneous answer recall.

### Degree-Weighted Path Counts as Metapath Features

We used the degree-weighted path count (DWPC) algorithm to quantify the connectivity between source and target nodes along a given metapath. This algorithm has been described extensively in other works^14,30^ and was recently used to produce machine learning features for the identification of novel Alzheimer’s disease genes.^31^ This algorithm can be described as follows:

### Degree-Weighted Path Counts (DWPC) Algorithm

#### Input

- *G = (V,E)* : A heterogeneous graph where *V* is the set of nodes and *E* is the set of edges.
- *M = (n*_*1*_,*e*_*1*_,*n*_*2*_,*e*_*2*_,*…,e*_*k*−*1*_,*n*_*k*_) : A metapath consisting of alternating node types *n*_*i*_ and edge types *e*_*i*_, representing a traversal pattern between different node types.
- *s ∈ V*_*s*_ *in V*_s_ *∈* V : Source node in the graph.
- *t ∈ V*_*t*_ *in V*_t_ *∈* V : Target node in the graph.
- *W* : Degree-weighting exponent, typically 0.0-1.0.

#### Output

- *DWPC(s,t,M)* : Degree-weighted path count between source node *s* and target node *t* along the metapath *M*.

1. **Initialization:**
  ∘ Let *P* be the set of all paths from *s* to *t* that conform to the metapath *M*. Each path *p ∈ P* will be a sequence of nodes and edges following the structure of *M*.
2. **Degree-Weighting:**
  ∘ For each path *p=(v*_*1*_,*e*_*1*_,*v*_*2*_,*…,e*_*k*−*1*_,*v*_*k*_*)∈P*, where *v*_*1*_*=* s and *v*_*k*_*=t*, calculate the degree-based weight as:

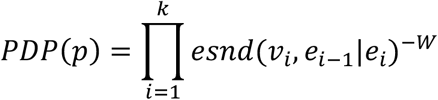

Here, *esnd*(*v*_*i*_, *e*_*i*−1_| *e*_*i*_) is the edge-specific node degree of *v*_*i*_ with respect to the edge type prior to *v*_*i*_ (*e*_*i*−1_) or after *v*_*i*_ (*e*_*i*_) in *M*. Thus, *v*_1_ and *v*_*k*_ will have only one *esnd* value while *v*_2_, *v*_3_, …, *v*_*k*−1_ will have two *esnd* values for linear *M*.
3. **Summing Over Paths:**
  ∘ The degree-weighted path count between *s* and *t* along the metapath *M* is the sum of the degree-weighted contributions from all paths in *P*:

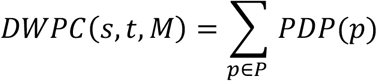
  ∘ If there are no valid paths between *s* and *t* that follow the metapath *M*, then *DWPC(s,t,M) = 0*.

For the datasets used in this work we selected an optimized dampening factor (*W*) of 0.2 in the DWPC algorithm. We then derived the DWPC as the summation of the PDP values across all answer paths, providing a comprehensive measure of connectivity that incorporates edge-specific node degree weighting.

### Target-Disease Scoring Model and Performance Evaluation

We produced an ensemble of 10 models using the scikit-learn Python package and the RandomForestClassifier method configured with 500 estimators, using the Gini Index to split the decision tree. These models were trained with input vectors of DWPC features for each Target-Disease node pair. Prior to training models, we held out an external dataset for model validation by selecting the target and disease nodes which only appear once in the set of positive pairs and removing any rows containing either of these target or disease nodes from the positive or negative training sets. The result is an external validation dataset with nodes entirely missing from the remaining training data, useful for testing the ability of the model to extrapolate to unseen data. For each model, we sampled all available positive training examples and randomly selected negative examples to create a 1:10 positive-to-negative ratio in the training set. This ratio was selected to reflect the massive imbalance of positive to negative pairs expected in a screening dataset generated during rule application. We performed a 5-fold cross-validation on each model to ensure reliable prediction accuracy on the internal training data.

To evaluate model performance on unseen data, we applied each model to predict outcomes on the full external dataset. We used the mean model prediction output among the 10 random forest classifiers as the final model prediction. We then compute AUROC, sensitivity (i.e., true positive rate), specificity (i.e., true negative rate), positive predictive value (PPV), and negative predictive value (NPV) for predictions on the external dataset. Then, to evaluate the stability of model prediction outputs, we calculated standard deviations of predictions across all 10 models on the external data. We then computed the mean value of standard deviations across all predictions to estimate model stability. Last, to evaluate utility of the ensemble model’s prediction output as a score for ranking hypothesis, we generated an early enrichment curve, which evaluates percentage recall of positive external data when rank-ordering all predictions by ensemble model score and checking the top *n* percent of predictions in the list.

### Retrieval-Augmented Generation (RAG) for Target Feasibility Reports

We generated large language model (LLM)-based target feasibility reports for top therapeutic target predictions to help guide the prediction review process. To create these reports, we first compiled a body of background knowledge and factual evidence by retrieving the top 10 rules, ranked by DWPC, for any given hypothesis. Then, we collected the first 100 answer paths, ranked by PDP, for each of these rules and transformed the answer paths into a natural language sentence (e.g., “*HIF1A* is target for cervical cancer, *FGFR3* is genetically associated with cervical cancer, and *FGFR3* is genetically associated with urinary bladder cancer.”). To refine the context, we computed text embeddings for each of these context sentences and calculated cosine similarities to a predefined prompt. The prompt posed the question:

> {question: “Is {target_name} a good therapeutic target for {disease_name}? Generate a detailed report on the feasibility of this target given what is known in the literature and the context above. Include details about genetic and molecular mechanisms relevant to the disease treatment.”}

Next, we selected the top 50 context sentences ranked by cosine similarity to the question prompt. Using these selected context sentences, we constructed the final LLM prompt, which included the following system and user instructions:

> {“role”: “system”, “content”: “You are a {disease_name} researcher searching for novel therapeutic targets to treat the disease. As an expert on {disease_name}, you should cite sources for your claims and provide references whenever possible.”}
>
> {“role”: “user”, “content”: “Given what you know from the {disease_name} literature and the following information: {context sentences}. Answer the question: {question}”}

The prompt was sent via the OpenAI chat completion API request with the GPT-4o model and temperature=0.7, and subsequently the target feasibility report was returned.

### Virtual Screening and Validation of Predicted Bladder Cancer Targets

We generated a DWPC feature dataset for bladder cancers by running ROBOKOP KG searches for all TEP rules with the specification that the DiseaseOrPhenotypicFeature node in the queries must be “urinary bladder cancer” (MONDO:0001187). In cases where specific rules yielded no results, the corresponding DWPC feature columns were populated with zeroes to maintain the dimensionality of the input dataset for model predictions.

To validate the predicted targets, we queried the ROBOKOP KG to identify drugs that physically interact with the target proteins. These drugs were either indicated for the treatment of urinary bladder cancer or had been involved in clinical trials for the disease. To retrieve this information, we employed the following Cypher queries:

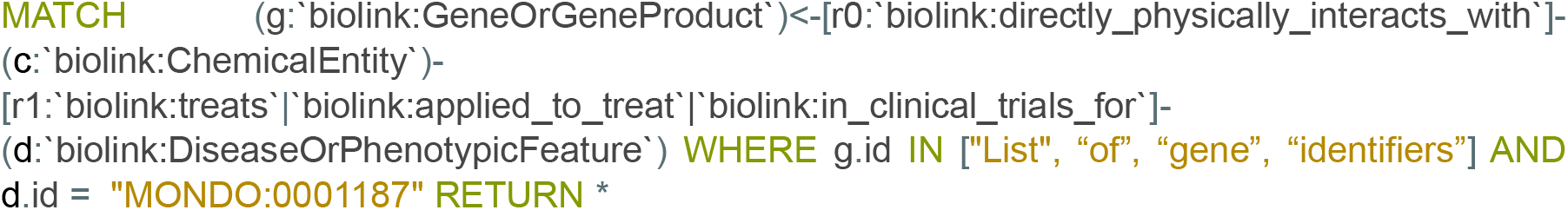

This query identified genes or gene products (g) that directly physically interact with chemical entities (c). These chemical entities are linked to disease or phenotypic features (d) through relationships indicating they treat, are applied to treat, or are involved in clinical trials for the disease. The condition d.id = “MONDO:0001187” ensures the query specifically focuses on urinary bladder cancer. Additionally, we applied the following query to capture activity-modulating interactions:

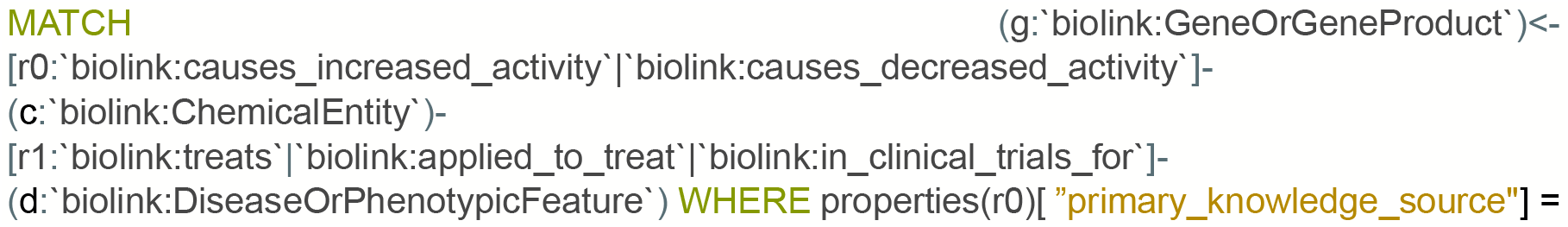
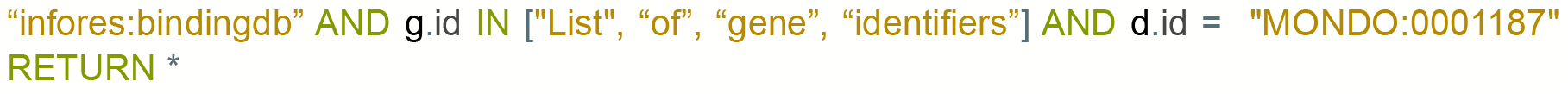

This query captured cases where chemical entities (c) are known to cause increased or decreased activity of gene products (g). These chemical entities are similarly associated with bladder cancer (d.id = “MONDO:0001187”) and are sourced explicitly from BindingDB (“primary_knowledge_source” = “infores:bindingdb”).^32^ Together, these queries enabled precise validation of target-drug interactions relevant to urinary bladder cancer, strengthening the confidence in our predicted therapeutic targets.

## Acknowledgements

The authors acknowledge ARPA-H Award Agreement # 140D042490001 and NIH (grant U24 GM146615) for partial funding of this research.

## Additional Information

### Supplementary Information

Supplementary information on target feasibility for cancer is available online.

### Data Availability

All data for this publication are available as supplementary information freely available with this paper.

