## Supplemental File 1 for "TARRAGON: Therapeutic Target Applicability Ranking and Retrieval-Augmented Generation Over Networks"

### Supplementary Note 1: RAG reports for target nomination

##### Feasibility of CDC6 as a Therapeutic Target for Urinary Bladder Cancer

**1. Introduction to CDC6:** Cell Division Cycle 6 (CDC6) is a key regulator in DNA replication and cell division, processes that are crucial for cancer progression. Its role involves the initiation of DNA replication by loading the MCM (minichromosome maintenance) complex onto DNA, thereby facilitating the unwinding and replication of the DNA strands. Overexpression or dysregulation of CDC6 has been associated with various cancers, including urinary bladder cancer, due to its role in promoting genomic instability and uncontrolled cellular proliferation.

**2. Genetic and Molecular Interactions:** The provided data underscores numerous interactions and regulatory mechanisms involving CDC6 that are pertinent to urinary bladder cancer:

- **CDC6 and TOP2A:** There is a consistent link between CDC6 and Topoisomerase II alpha (TOP2A), a known target in urinary bladder cancer. CDC6 interacts with multiple proteins (e.g., CDK1, CDK4, CDK6, MYC) that regulate or genetically interact with TOP2A. Given TOP2A's role in DNA replication and its targeting by existing chemotherapies (e.g., etoposide, doxorubicin), CDC6's interaction with TOP2A pathways suggests that CDC6 influences TOP2A activity indirectly, making CDC6 a potential upstream target.
- **Cell Cycle Regulation:** CDC6 interacts with cyclin-dependent kinases (CDK1, CDK2, CDK4, CDK6) and cyclins (e.g., CCNB1), which are critical for cell cycle progression. Dysregulation of these interactions can lead to increased cell proliferation, a hallmark of cancer. Inhibiting CDC6 could disrupt this regulatory network, potentially halting tumor growth.
- **Interaction with Tumor Suppressors and Oncogenes:** CDC6 interacts with CDKN1A (p21), a cyclin-dependent kinase inhibitor that regulates cell cycle progression. Modulation of CDC6 may affect the tumor suppressive functions of CDKN1A. Additionally, CDC6's interaction with MYC, a well-known oncogene, further implicates it in oncogenic pathways.
- **DNA Replication and Repair:** CDC6 is involved in DNA replication initiation. Its interaction with MCM proteins and PCNA, both involved in DNA replication and repair, suggests that targeting CDC6 could impair the DNA replication machinery, leading to replication stress and cell death in cancer cells.

**3. Potential Therapeutic Agents:** Several drugs currently in clinical trials or use for bladder cancer treatment have been identified to interact with pathways involving CDC6:

- **Docetaxel and Paclitaxel:** These taxanes affect microtubule dynamics and have been shown to downregulate proteins interacting with CDC6.

- **Gefitinib and Sunitinib:** These tyrosine kinase inhibitors have demonstrated effects on CDC6-related pathways.
- **Gemcitabine and Mitomycin:** Chemotherapeutic agents that upregulate or influence pathways involving CDC6.
- **Bortezomib and Irinotecan:** Both have shown regulatory effects on proteins interacting with CDC6, such as CDKN1A.

**4. Challenges and Considerations:**

- **Target Specificity:** Given CDC6's essential role in cell division, systemic inhibition might affect normal proliferating cells leading to toxicity. Strategies to selectively target cancer cells, such as conjugating CDC6 inhibitors with tumor-specific antibodies or using nanoparticle delivery systems, could mitigate this risk.
- **Resistance Mechanisms:** Cancer cells might develop resistance to CDC6 inhibition. Combination therapies targeting multiple pathways could help overcome resistance.

**5. Conclusion:** CDC6 is a promising therapeutic target for urinary bladder cancer due to its central role in DNA replication and cell cycle regulation, and its interaction with oncogenic and tumor suppressive pathways. While further preclinical and clinical investigations are necessary, the existing data supports the potential of CDC6-targeted therapies, either alone or in combination with current chemotherapeutic agents, to enhance treatment efficacy for urinary bladder cancer.


#### MDM2 as a Therapeutic Target for Urinary Bladder Cancer: A Detailed Report

##### Introduction

MDM2 (Mouse Double Minute 2 homolog) is an oncogene that encodes a protein functioning as a critical negative regulator of the tumor suppressor p53. It is known to be overexpressed in various cancers, leading to the degradation of p53, thereby promoting tumorigenesis. MDM2 has been genetically associated with urinary bladder cancer, suggesting its potential as a therapeutic target.

##### Genetic Association and Mechanism

- **MDM2 Genetic Association with Urinary Bladder Cancer:** MDM2 is frequently amplified in urinary bladder cancer, suggesting its role in tumor progression and maintenance. This makes MDM2 a plausible target for therapeutic intervention.
- **Interplay with p53:** MDM2 directly interacts with p53, leading to its ubiquitination and subsequent proteasomal degradation. Inhibition of MDM2 can restore p53 function, leading to cell cycle arrest and apoptosis in cancer cells with a functional p53 pathway.

- **Interaction with TOP2A:** MDM2 interacts with several proteins that regulate TOP2A, a well-known target in bladder cancer therapy. These interactions suggest that MDM2 might play a role in modulating the efficacy of TOP2A-targeting drugs, potentially enhancing their therapeutic effects.
- **Other Genetic Interactions:** MDM2 interacts with numerous proteins and pathways involved in cancer progression, such as UBC, UBE2D1, USP7, and TP53, indicating its central role in cellular processes relevant to bladder cancer.

**Feasibility as a Therapeutic Target**

- Targeting MDM2 in p53 Wild-Type Bladder Cancers: In tumors where p53 is not mutated, MDM2 inhibition could potentially restore p53's tumor-suppressive functions.
- Combination Therapies: MDM2 inhibitors could be combined with chemotherapeutics targeting TOP2A or other pathways to enhance antitumor efficacy.

**Candidate Drugs and Interacting Compounds**

- **Nutlin-3:** A well-known small molecule inhibitor of MDM2-p53 interaction. It has shown promise in preclinical studies for various cancers by activating p53-dependent transcription and inducing apoptosis.
- **MI-773:** Another MDM2 antagonist that has shown potential in restoring p53 function and inhibiting tumor growth in preclinical models.
- **Idasanutlin (RG7388):** An orally available MDM2 inhibitor in clinical trials for several cancers. It could be a candidate for trials in urinary bladder cancer.
- **Doxorubicin and Gemcitabine:** While not direct MDM2 inhibitors, they have shown interactions with the MDM2 pathway or its downstream effects, suggesting combinatorial potential with MDM2-targeting drugs.
- **Bortezomib:** Known to modulate proteins interacting with MDM2, suggesting a potential synergistic effect when combined with MDM2 inhibitors.

**Challenges and Considerations**

- Selective Targeting: As MDM2 is involved in regulating p53, care must be taken to avoid adverse effects due to excessive p53 activation, which could lead to unwanted cell death in normal tissues.
- Resistance Mechanisms: Tumors with mutant p53 may not respond to MDM2 inhibition, necessitating the stratification of patients based on p53 status.
- Clinical Development: While preclinical data is promising, more clinical trials are needed to establish the safety and efficacy of MDM2 inhibitors in urinary bladder cancer.

**Conclusion**

MDM2 is a promising therapeutic target for urinary bladder cancer, especially in tumors with intact p53. The genetic and molecular data support the potential for MDM2 inhibitors to be integrated into treatment regimens. Given the complexity of cancer pathways and potential for drug resistance, combination therapies and patient stratification will be critical for the successful clinical application of MDM2-targeting strategies. Further research and clinical trials are warranted to validate these findings and optimize therapeutic approaches.

**Report: Evaluating GART as a Therapeutic Target for Urinary Bladder Cancer**

**Introduction**

Urinary bladder cancer (UBC) is a prevalent malignancy with significant morbidity and mortality. Identifying novel therapeutic targets is crucial to improving treatment outcomes. This report evaluates the feasibility of targeting GART (glycinamide ribonucleotide formyltransferase) in UBC, leveraging its known associations and interactions in other cancers, particularly non-small cell lung carcinoma (NSCLC).

**GART Overview**

GART is an enzyme involved in the purine biosynthesis pathway, crucial for nucleotide production. Its role in cancer involves facilitating rapid cell proliferation by providing necessary nucleotide resources. Targeting metabolic pathways in cancer cells is a promising strategy, as these pathways are often upregulated to meet the demands of rapidly dividing tumor cells.

**Evidence Supporting GART as a Target in Cancer**

**1. GART in Non-Small Cell Lung Carcinoma (NSCLC)**

- **Target Role:** GART has been identified as a therapeutic target in NSCLC, suggesting its critical role in tumor metabolism and growth.

- **Drug Interaction:** Potential inhibitors of GART have been explored in NSCLC, indicating its druggability.

##### 2. Genetic and Molecular Associations

- **GART and Disease Models:** GART is genetically associated with various disease models, indicating a fundamental role in disease progression, including cancer.
- **Genetic Interactions:** GART interacts with other proteins like UBC and CUL5, which are involved in cancer-related pathways. UBC regulates TOP2A, a known target in UBC, suggesting potential pathway overlaps.

##### 3. Potential in Urinary Bladder Cancer

While direct evidence of GART's role in UBC is limited, its involvement in similar metabolic pathways in NSCLC supports its consideration as a target. The metabolic demands of UBC may be similar, given the tumor's rapid growth characteristics.

##### Candidate Drugs Interacting with GART

Given GART's role in NSCLC, several drugs and compounds that interact with GART or its associated pathways could be repurposed or further investigated for UBC:

- **Methotrexate:** Although primarily known for inhibiting dihydrofolate reductase, it affects the folate pathway, indirectly impacting GART's activity. Methotrexate is already used in UBC, providing a potential combinatorial approach.
- **Gemcitabine:** Commonly used in UBC, it targets DNA synthesis. Combining it with GART inhibitors could enhance anti-tumor efficacy by disrupting nucleotide synthesis.
- **Fluorouracil:** Known to interfere with nucleotide synthesis, it could act synergistically with GART inhibition.

##### Feasibility and Challenges

##### Feasibility

- **Metabolic Targeting:** Given the success of metabolic targeting in other cancers, exploring GART in UBC is promising. Its role in nucleotide biosynthesis is crucial for tumor growth.
- **Pathway Interactions:** The genetic interactions of GART with known UBC targets like TOP2A provide a rationale for its exploration.

##### Challenges

- **Specificity:** Ensuring that GART inhibitors selectively target cancer cells without affecting normal proliferating tissues is crucial to minimize side effects.
- **Resistance Mechanisms:** Like other metabolic targets, resistance mechanisms could develop, necessitating combination therapies.

#### Conclusion

GART represents a novel and potentially effective therapeutic target for urinary bladder cancer, especially given its established role in NSCLC and the metabolic demands of cancer cells. The feasibility of targeting GART lies in its metabolic function and interaction with cancer-related pathways. Further preclinical studies are necessary to elucidate its specific role in UBC and to identify potent inhibitors for clinical development.

#### References


- Comprehensive studies on GART's role in NSCLC and its potential inhibitors.
- Current literature on metabolic targeting in cancer therapy.
- Existing clinical trials involving GART-related pathways and drugs in various cancers, including UBC.


This report highlights the potential of GART as a targeted therapy in UBC and calls for further research to validate these findings and develop effective therapeutic strategies.

#### Feasibility of LLPH as a Therapeutic Target for Urinary Bladder Cancer

#### Introduction

Urinary bladder cancer is a prevalent malignancy with a need for novel therapeutic targets to improve treatment outcomes. In this context, LLPH (LLP homolog) emerges as a potential target due to its interactions with known cancer-related proteins and its involvement in cellular mechanisms pertinent to cancer biology.

#### Genetic and Molecular Mechanisms

1. **Interaction with TOP2A**:
    a. LLPH physically interacts with TOP2A, a well-established target in urinary bladder cancer. This interaction occurs both directly and through a network of associated proteins, indicating a potential role in the regulation of TOP2A activity. TOP2A is essential for DNA replication and cell proliferation, processes frequently dysregulated in cancer (Wang, 1996).
2. **Co-expression with Cancer-related Genes**:
    a. LLPH co-expresses with multiple genes such as PHF5A, HMGA2, and NCAPG, which regulate or affect the response to chemotherapeutic agents like Mitomycin. This co-expression suggests LLPH's involvement in pathways that modulate drug sensitivity and cancer cell survival.
3. **Regulation by Chemotherapeutic Agents**:
    a. Several known chemotherapeutic agents, including Gemcitabine, Doxorubicin, and Mitomycin, modulate the expression of proteins interacting with LLPH, such as CTCF and TOE1. This regulation implies that

LLPH may serve as a mediator of chemotherapeutic efficacy, potentially enhancing the therapeutic response when targeted.

4. **Interaction with Ribosomal Proteins**:
    a. LLPH interacts with various ribosomal proteins (e.g., RPL4, RPL8, RPL36), which are essential for protein synthesis and cellular growth. Alterations in ribosomal biogenesis and function are a hallmark of cancer (Pelletier et al., 2018), suggesting that LLPH may influence cancer cell metabolism and proliferation.
5. **Involvement in Nucleolar Function**:
    a. LLPH is active in the nucleolus, a site of ribosome biogenesis and a stress sensor in cells. Its role in nucleolar function underscores its potential impact on cellular stress responses, which are critical in cancer development and progression.

**Therapeutic Potential and Candidate Drugs**

1. **Targeting LLPH Directly**:
    a. Developing small molecules or biologics that specifically inhibit LLPH function could disrupt its interactions with critical oncogenic proteins, such as TOP2A, potentially leading to reduced tumor growth and enhanced sensitivity to existing treatments.
2. **Candidate Drugs**:
    a. **Mitomycin**: Altering LLPH activity might enhance the response to Mitomycin, given the co-expression and regulatory interactions with HMGA2 and other associated proteins.
    b. **Gemcitabine and Doxorubicin**: These drugs modulate proteins interacting with LLPH, suggesting that LLPH inhibition might synergize with these agents to improve therapeutic outcomes.
3. **Considerations for Drug Development**:
    a. The development of LLPH-targeting drugs should focus on disrupting its interaction networks, especially those involving TOP2A and ribosomal proteins. Additionally, considering its nucleolar localization, targeting LLPH could also affect cellular processes related to ribosome biogenesis and stress responses.

**Conclusion**

LLPH presents as a promising therapeutic target in urinary bladder cancer due to its extensive interaction network with cancer-associated proteins and its involvement in crucial cellular processes such as DNA replication, protein synthesis, and stress response. Further preclinical studies are necessary to elucidate the precise role of LLPH in bladder cancer pathophysiology and to validate its potential as a therapeutic target. The integration of LLPH-targeting strategies with existing chemotherapies could pave the way for more effective treatments.
